# A lightweight deep-learning detector for the real-time monitoring of the invasive frogs *Rhinella marina* and *Polypedates leucomystax*

**DOI:** 10.64898/2026.10.05.756611

**Authors:** Kosmas Deligkaris, Melika Sadeghi Tabrizi, Greg Stephens

## Abstract

Early detection of invasive species is critical to efficiently managing biological invasions. Passive acoustic monitoring combined with deep learning has become an effective tool for identifying invasive anurans from their species-specific mating calls. In practice, however, these systems still depend on manually retrieving recordings and running inference on a dedicated workstation, which delays detection at invasion fronts, especially on remote islands where frequent travel is impractical. Here, we present a lightweight convolutional neural network for detecting *Rhinella marina* and *Polypedates leucomystax* that can be used with low-power microcontrollers, enabling continuous, real-time monitoring of these invasive species. Our model achieved a mean invasive-species ***F***_**1**_ of 0.87 with precision higher than 0.88 for both species, comparable to the performance of substantially larger models at a fraction of the model size. Cross-domain evaluation confirmed reliable detection on two out-of-training locations: Iriomote Island, Japan, and on an independent dataset from Australia, demonstrating generalization across recording conditions and biogeographic contexts, albeit with local threshold calibration for *P. leucomystax* at Iriomote. Our results demonstrate that effective acoustic detection of invasive species is feasible on self-contained, battery-powered devices, removing the dependence on server infrastructure and enabling autonomous early warning systems in remote protected areas.

## 1 Introduction

Biological invasions are a leading driver of biodiversity loss and ecosystem degradation. The reported global damage and management costs exceed US$1.2 trillion over recent decades and continue to rise steeply [1]. As eradication is only feasible early in an invasion, while a population is still small and localized, early detection is paramount. Once an invasive species becomes established, eradication is often impractical, and only costly, long-term management remains [2]. Nonetheless, management remains largely reactive, with action taken on average 11 years after a species’ impacts are first reported, while spending on prevention is roughly 25 times lower than on post-establishment control [3]. Early-warning systems and technologies that detect new invasions rapidly can be fundamental to shifting management of invasive species from reactive to proactive, supporting early detection and rapid response (EDRR) while eradication remains feasible [4].

The Ryukyu Archipelago in southwestern Japan hosts globally significant endemic diversity, with parts of the archipelago (e.g., Iriomote, Amami-Oshima, Tokunoshima) designated a UNESCO Natural World Heritage Site. The globally endangered frogs *Odorrana splendida* [5], *Odorrana supranarina* [6], and *Odorrana utsunomiyaorum* [7] are all endemic to these islands. The region’s vertebrate fauna is similarly rich in endemic, threatened species, including the Iriomote cat (*Prionailurus bengalensis iriomotensis*) [8] and the Ryukyu crested serpent eagle (*Spilornis cheela perplexus*) [9]. As these species are found nowhere else and often exist in small numbers, they are especially vulnerable to invasive species.

Ishigaki island, which is only 30 km from Iriomote, hosts established populations of the two invasive species *R. marina* and *P. leucomystax. R. marina* has been identified repeatedly in Iriomote in areas near the port, and was most likely transported from Ishigaki with cargo and construction materials [10]. Control programs have so far kept these species from establishing in Iriomote, but they rely primarily on opportunistic observations by local citizens [10]. As neither species is yet established in Iriomote, early detection of these invasive species can enable eradication before populations establish.

The cane toad (*Rhinella marina*) is native to the Americas and listed among the 100 of the World’s Worst Invasive Alien Species. It is best known from its introduction to Australia in 1935, where its toxic secretions have caused severe declines in native predators that attempt to consume it [11]. Population collapses exceeding 80% have been documented in large-bodied predators such as monitor lizards, elapid snakes, freshwater crocodiles, and dasyurid marsupials [11, 12]. Toads continue to expand into new regions where further biodiversity losses are predicted [13].

The Southeast Asian treefrog (*Polypedates leucomystax*) was first recorded in the Ryukyu Archipelago in 1964 and has since established feral populations on more than 36 islands [14, 15]. An opportunistic feeder with a broad arthropod diet [15], it competes with native anurans for food and breeding sites and can introduce parasites to native amphibian communities [16]. The continued spread of both species toward currently uninvaded islands, including protected areas, makes their early detection at invasion fronts a conservation priority.

Because male anurans produce species-specific calls during breeding, passive acoustic monitoring (PAM) combined with deep learning has made automated detection of anuran invasive species feasible [17]. Kimura et al. [18] demonstrated this for *P. leucomystax* and *R. marina*, training a BirdNET-based detector that achieved high precision and recall on recordings from the Ryukyu Archipelago. Similarly, Bota et al. [19] developed a BirdNET-based detector of the American bullfrog with a precision as high as 99.7%. BirdNET is a widely used convolutional neural network model for bird (and increasingly other taxa) species identification, trained on large annotated acoustic datasets [20].

A major disadvantage of conventional PAM devices is that they store audio on SD cards, which must be retrieved manually before analysis can take place. On remote islands, where cost and logistics make frequent visits impractical, this introduces delays of weeks to months between an invasion and its detection. During this time, an invasive population may establish and spread widely, substantially increasing management costs and reducing the probability of eradication [21].

An alternative approach is the use of Edge AI, in which a detection model runs directly on the recording device. Such devices can analyze the soundscape in real time and, combined with low-bandwidth wireless links, report detections to eradication teams within hours rather than months. Several systems have moved acoustic classification toward the edge, but all rely on comparatively powerful single-board computers. BirdNET-Pi (https://github.com/mcguirepr89/BirdNET-Pi) runs a TFLite build of BirdNET on a Raspberry Pi, while the Bird@Edge system [22] runs an EfficientNet-B3 model on an NVIDIA Jetson Nano. Such single-board computers draw several watts of power and depend on substantial battery and solar energy for continuous operation.

By contrast, a microcontroller unit (MCU) draws substantially lower power than a single-board computer, and is designed for long-term, low-maintenance operation in embedded systems. A single MCU-based device could therefore record, classify, and report detections autonomously for extended periods on a small battery and modest solar panel, and could be left running at remote sites that are hard to reach and service. Utilization of MCU-based Edge AI sensors enables a monitoring strategy that complements larger, power-hungry stations based on single-board computers: rather than a few central installations, multiple small, distributed sensors can be placed across a variety of locations, an approach that is well matched to invasive species such as *P. leucomystax* that breed and disperse across many small, scattered sites. Unfortunately, as MCUs have substantial compute limitations (memory, processing power) they also require a significantly smaller deep-learning model that may not match the detection performance of the larger architectures.

Here, we present a lightweight deep-learning detector for *P. leucomystax* and *R. marina* designed for low-cost, battery-powered MCU devices. Trained on the dataset of Kimura et al. [18], our model contains approximately 33,000 parameters and occupies 67 KB once quantized, which is small enough to run on MCU-class hardware, enabling a mode of environmental monitoring that is not currently feasible with single-board computers.

## 2 Materials and Methods

### 2.1 Datasets

In this study, we utilized two datasets. The primary dataset, used for both training and evaluation, was that of Kimura et al. [18]. Briefly, the dataset contains weakly labeled 3-second audio clips of eight anuran species, including the two invasive species of interest, *P. leucomystax* and *R. marina*. Recordings were collected at two locations in the Ryukyu Archipelago: Ishigaki island and Iriomote island. Recordings from Ishigaki Island, where both invasive species are established, were used for training, validation, and testing. As Iriomote island does not host these invasive species, recordings were conducted through a playback operation, where the calls of the two invasive species were broadcasted and re-recorded in the field (see [18] for full details).

To further evaluate geographic generalization, we used the Australian cane toad dataset [23], which contains weakly labeled 3-second clips across 11 classes, including *R. marina*, background noise, and other species, recorded by the Australian Acoustic Observatory across a range of locations and environmental conditions [24]. Since *R. marina* is common to both datasets, we used the Australian recordings as a second evaluation dataset, testing whether our detector generalizes across continents without any additional training. For this evaluation, because all other species differ between the two datasets, we grouped them into a single category, “others” and focused only on the detection of *R. marina*. Weakly labeled samples are defined as 3 s samples that include the label of the species present, but not its exact timing within this period.

### 2.2 Audio preprocessing

All audio was resampled to 16 kHz and low-pass filtered at 6 kHz to suppress out-of-band insect noise. Each clip was converted to a log-mel spectrogram using a short-time Fourier transform with an FFT window of 1024 samples (64 ms) and a hop length of 750 samples (47 ms). The power spectrum was projected onto 128 mel bands. A 3-seconds clip produces 64 time frames, resulting in a (128 x 64) single-channel input to our lightweight CNN detector. The same preprocessing was applied across all datasets and for all comparison architectures.

### 2.3 Pipeline and CNN model architecture

The lightweight detector is a compact CNN that takes a (128 x 64) log-mel spectrogram as input and produces eight independent sigmoid outputs, one per species. The sigmoid output of each species is then thresholded independently to produce a binary detection result for each of the eight species (Figure 1a).

**Fig. 1.**
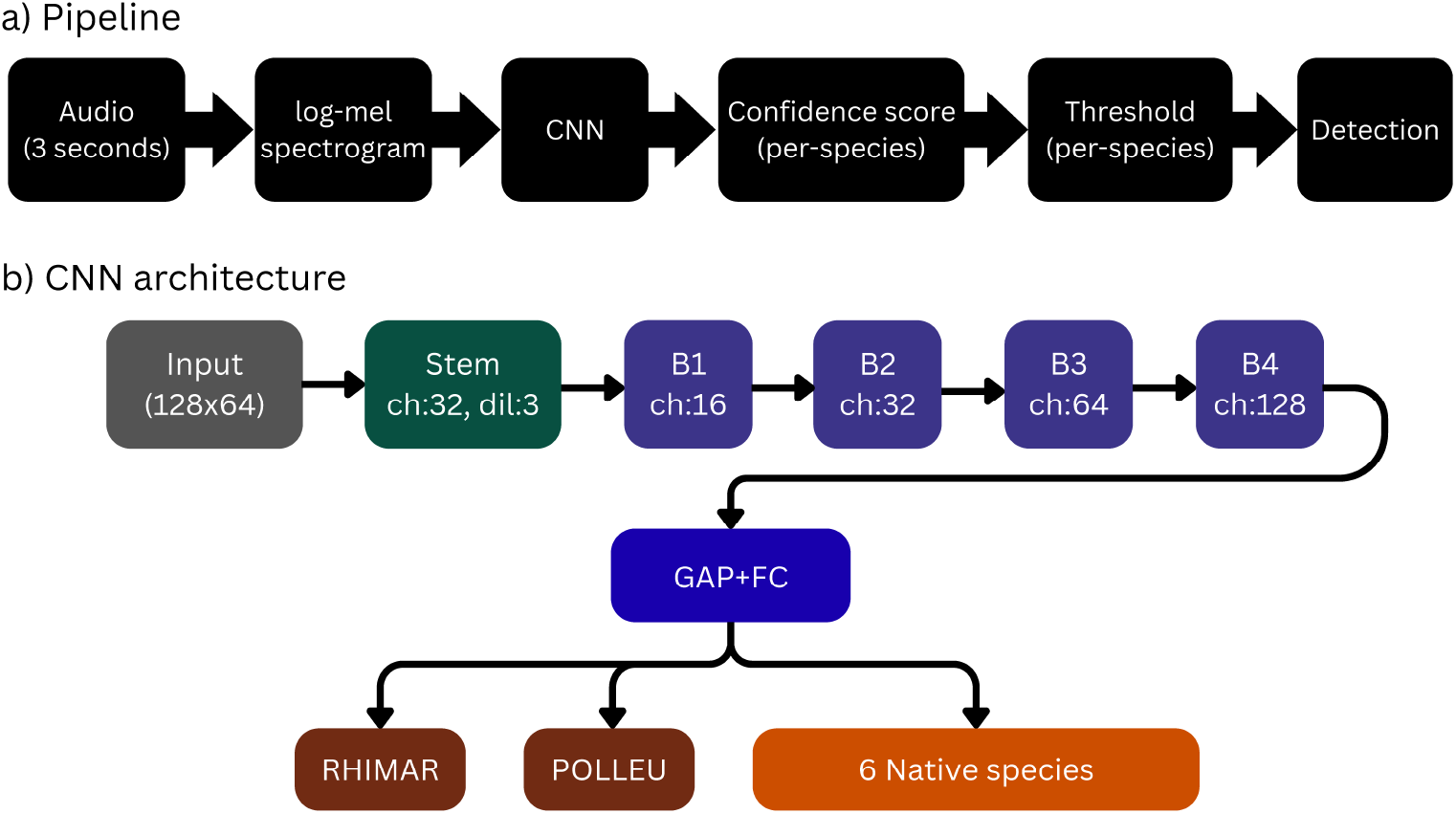
a) Overall pipeline for detecting and classifying anuran species. b) Architecture of the lightweight CNN model. Our lightweight detector is composed of a dilated stem block, followed by four consecutive blocks (B1,…,B4), global average pooling (GAP) and a fully connected layer (FC). Each block is composed of a series of depthwise convolution, pointwise convolution, a squeeze and excitation layer and a residual connection followed by 2x2 max-pooling. The final output of the model is eight sigmoid outputs for the eight species present in the training dataset (RHIMAR, POLLEU as the invasive species plus six native species)

**Fig. 2.**
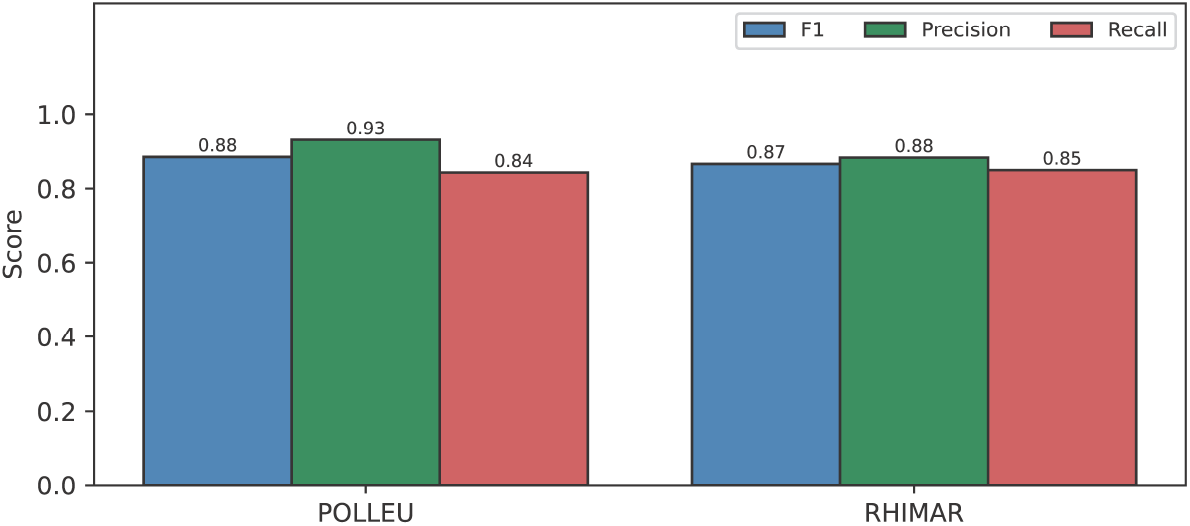
Performance of the lightweight model in wild sound recordings in Ishigaki island. POLLEU: *P. leucomystax*, RHIMAR: *R. marina*

The CNN architecture combines depthwise separable convolutions [25] with squeeze-and-excitation blocks [26], both widely used for resource-constrained models. The network begins with a convolutional stem: a 32-filter 3x3 convolution with dilation rate 3, followed by batch normalization, a ReLU activation, and 2x2 max-pooling. The dilated stem widens the receptive field over the input spectrogram before the first downsampling step. The stem feeds four sequential blocks, each consisting of a 3x3 depthwise convolution, a 1x1 point-wise convolution (each followed by batch normalization and ReLU), a squeeze-and-excitation module that reweights channels by learned importance (reduction ratio 8), and a residual connection across the block. A 1x1 projection is applied on the residual path where the channel count changes. Each block is followed by 2x2 max-pooling, halving the spatial resolution. The four blocks have output channel widths of 16, 32, 64, and 128, so the feature maps progress from 128 x 64 at the input to 4 x 2 after the final block. A global average pooling layer then feeds a fully connected layer with eight sigmoid outputs (see Figure 1b). Training the full eight species head allows the native species to serve as discriminative negatives for the two invasive targets. The complete model contains 33,208 parameters and occupies 66.6 KB after INT8 quantization, small enough for deployment on low-power microcontrollers.

### 2.4 Training procedure

The model produces eight independent sigmoid outputs, one per species, and was trained on all eight species simultaneously as a multi-label task. This design lets the six native species act as discriminative negatives for the two invasive targets: by learning to recognize the calls that co-occur in the same soundscape, the model can better distinguish *P. leucomystax* and *R. marina* from acoustically similar native species.

To prevent the model from neglecting under-represented classes, we trained with an asymmetric loss function (ASL [27]), which penalizes missed detections (false negatives) more heavily than false positives. To further counteract the class imbalance, each class’s contribution to the loss was weighted by the inverse of its effective number of samples [28], up-weighting the rare classes’ contribution relative to the common ones.

The model was trained for up to 150 epochs using the Adam optimizer (learning rate 10^−3^, batch size 32) on an 85/15 training/validation split of the Ishigaki training data. The learning rate was reduced on validation-loss plateaus, and training was stopped early when validation loss ceased to improve (patience: 25 epochs), restoring the best-performing weights. To improve robustness to the acoustic variation encountered across recording sites, we applied a two-stage augmentation pipeline during training: time shifting, gain variation, additive Gaussian and pink noise, pitch shifting, and time stretching were applied to the raw audio, and SpecAugment time- and frequency-masking was applied to the resulting spectrogram [29]. Validation and test data were not augmented.

After training, the model was converted to an INT8 TFLite model using post-training quantization. This quantized model is the one that can be deployed on-device and used to compute all the reported metrics.

### 2.5 Comparison architectures

To assess how our lightweight detector compares against established, larger architectures, we trained three larger convolutional networks on the same task. Specifically, we trained EfficientNet-B0 [30], EfficientNet-B0 at half width, and ResNet50 pretrained on ImageNet [31]. Parameter counts and model sizes are reported in Table 6.

All baselines used the same preprocessing, training pipeline, augmentation, loss, and threshold-selection procedure as our lightweight model. Each model was calibrated identically: per-species *F*_1_-optimal thresholds were derived from the validation split and subsequently applied unchanged to the test data.

### 2.6 Evaluation protocol and threshold selection

For each species, detection is treated as a binary classification problem over the 3-second windows samples of the test dataset, and performance is quantified using precision, recall, and the *F*_1_ score. Denoting by *TP, FP*, and *FN* the numbers of true positives, false positives, and false negatives, respectively, precision (*P* ) and recall (*R*) are defined as:

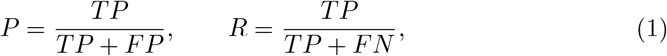

where precision quantifies the proportion of positive predictions that are correct and recall the proportion of true positives that are recovered. Practically, a lower recall would increase the probability of missing an invasive species, while lower precision would increase the “false alarms” of the detector. The *F*_1_ score is the harmonic mean of precision and recall, defined as:

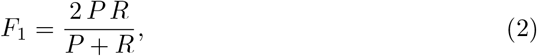

The *F*_1_ score provides a single summary statistic that is maximized only when both precision and recall are jointly high.

The detector’s output is a confidence score for each independent class (species), ranging from 0 to 1, which is then thresholded to produce a single True/False value for each class. The selection of the threshold can be different for each class, allowing practitioners to establish high-performing models tailored to the properties of the local soundscape (species presence, background noise, etc).

For the Ishigaki test set, we selected a per-species threshold that maximized the *F*_1_ score on the validation set, sweeping a grid of 91 candidate values evenly spaced from 0.05 to 0.95, and applied this threshold unchanged to the held-out test set. Selecting the threshold on validation data rather than on the test set itself is essential for a realistic performance estimate, as a detector deployed in the field would not have access to labeled data (represented by the test set).

For the Iriomote and Australia test sets, we evaluated our detector with two different thresholds. The first threshold used was the one found from the Ishigaki validation set, applied unchanged to the Iriomote and Australia test sets. This threshold would represent a “zero-shot” evaluation of our detector on these datasets. The second threshold was found by conducting the same sweep as for the Ishigaki dataset, but this time directly on each of the two test sets. Using this threshold allowed us to characterize the capacity of our detector without the confounding factor of threshold selection. In practice, utilizing this approach would necessitate local recordings, either from real frogs or from playback recordings (as in the case of e.g., Iriomote island, as analyzed here and in [18]).

### 2.7 Ablation experiments

To verify that our detector’s performance arises from features learned during training rather than from the architecture or threshold alone, we evaluated two ablated variants against the fully trained model. In the first variant, the convolutional backbone was simply set to random weight values, without any training. Only the final, fully connected classification layer was trained, testing how well a trained classifier performs on top of random convolutional features. In the second variant, both the convolutional backbone and the classification layer were set at random values, providing a lower-bound estimate of our detector’s performance attributable to the architecture and threshold, in the absence of any learning. Both variants used the same architecture, training data, and evaluation procedure as the full model.

### 2.8 Microcontroller benchmarking

To determine which MCU can run our detector in real-time, an estimate of its performance in various MCU and overall memory requirements is needed. Three key metrics were used to assess device compatibility: Random Access Memory (RAM), Read-only Memory (ROM), and inference time. For a deep learning model, ROM stores the model’s weights, while RAM stores temporary activations during inference. Lastly, inference time is the time required to run the model and complete inference on a 3 s spectrogram. On MCUs, the most restrictive factor is usually the peak RAM needed to hold a model’s intermediate activations, rather than the flash needed to store the model itself [32].

We obtained these estimates using Edge Impulse [33], a platform for developing and profiling machine-learning models on embedded hardware. Given an exported model, Edge Impulse compiles it and can provide estimates of the inference latency, RAM, and ROM memory requirements. We profiled the INT8 model across a range of MCU platforms, ranging from highly resource-constrained devices to more capable boards with dedicated neural-processing hardware. The evaluated platforms are summarized in Table 1 and described briefly below.

**Table 1.**
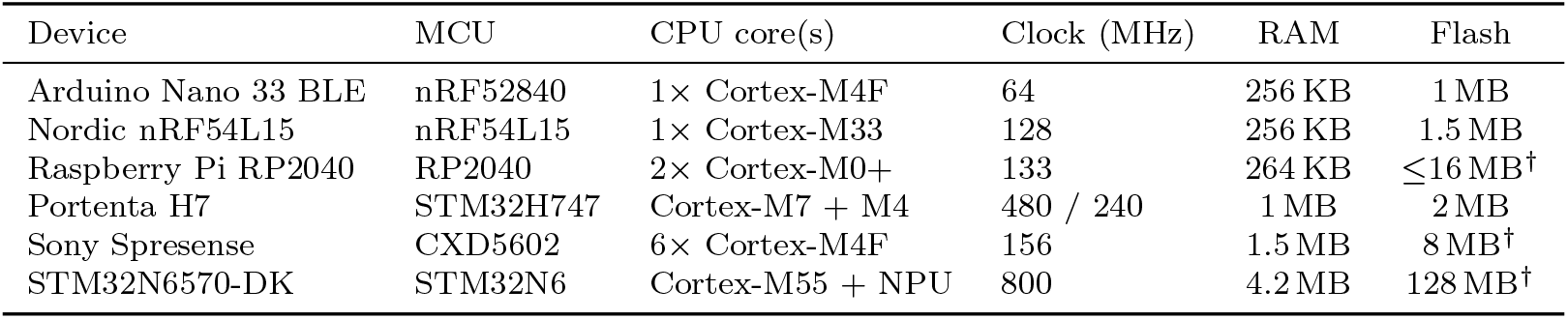
Specifications of the profiled microcontroller platforms (ordered by on-chip RAM). The platforms span a broad range of low-power compute, from single-core Cortex-M0+/M4F devices to a Cortex-M55 with an integrated Neural-ART NPU. RAM refers to on-chip SRAM, used for writing the model activations, while flash is the program-storage capacity, also used for storing the model weights. ^*†*^Flash is external to the MCU; the value shown is the board/kit’s flash device.

| Device | MCU | CPU core(s) | Clock (MHz) | RAM | Flash |
| --- | --- | --- | --- | --- | --- |
| Arduino Nano 33 BLE | nRF52840 | 1× Cortex-M4F | 64 | 256 KB | 1 MB |
| Nordic nRF54L15 | nRF54L15 | 1× Cortex-M33 | 128 | 256 KB | 1.5 MB |
| Raspberry Pi RP2040 | RP2040 | 2× Cortex-M0+ | 133 | 264 KB | ≤16 MB<sup>†</sup> |
| Portenta H7 | STM32H747 | Cortex-M7 + M4 | 480 / 240 | 1 MB | 2 MB |
| Sony Spresense | CXD5602 | 6× Cortex-M4F | 156 | 1.5 MB | 8 MB<sup>†</sup> |
| STM32N6570-DK | STM32N6 | Cortex-M55 + NPU | 800 | 4.2 MB | 128 MB<sup>†</sup> |

The Arduino Nano 33 BLE is the most restrictive of the evaluated platforms, containing only 256 KB of RAM and 1 MB ROM, based on the nRF52840 MCU from Nordic Semiconductors, running at 64 MHz. The Nordic nRF54L15 runs at 128 MHz, with 256 KB RAM and 1.5 MB Flash memory. The Raspberry Pi RP2040 uses Arm’s dual Cortex-M0 at 133 MHz, with 264 KB RAM and up to 16 MB external Flash memory. The Arduino Portenta H7 is a higher-spec MCU with a dual-core (Cortex-M7/M4) running at 480 MHz and 240 MHz, respectively with 1 MB RAM and 2 MB Flash memory. The Sony Spresense is an ultra-low power, multi-core MCU with six Cortex-M4F at 156 MHz, with 1.5 MB RAM and 8 MB Flash memory. Lastly, the STM32N6570-DK (development kit) utilizes the STM32N6 chip, featuring a Cortex-M55, along with a Neural Processing Unit (NPU) that can accelerate deep learning computations. It runs at 800 MHz and has 4.2 MB RAM and 128 MB Flash memory.

### 2.9 LLM use

Part of the code used in this study was written with the assistance of a large language model (Claude, Anthropic). All code was reviewed by the authors, who take full responsibility for its correctness and for the results reported.

## 3 Results

### 3.1 Performance on wild sounds in Ishigaki island

Our lightweight detector achieved an *F*_1_ score of 0.88 for *P. leucomystax* and 0.87 for *R. marina*, with high precision for both species (0.93 and 0.88, respectively). The optimal threshold for the two invasive species of interest was 0.62 for *P. leucomystax* and 0.79 for *R. marina*. These are the thresholds that maximize the *F*_1_ score for each invasive species in its respective validation set (Table 2).

**Table 2.** Per-species detection on the held-out wild-sound test set recorded at Ishigaki, for the two invasive species POLLEU (*P. leucomystax*) and RHIMAR (*R. marina*). TP/FP/FN = counts of true positives, false positives, and false negatives over 3-second windows.

| Species | TP | FP | FN | F1 | Precision | Recall | Threshold |
| --- | --- | --- | --- | --- | --- | --- | --- |
| POLLEU | 107 | 8 | 20 | 0.88 | 0.93 | 0.84 | 0.62 |
| RHIMAR | 45 | 6 | 8 | 0.87 | 0.88 | 0.85 | 0.79 |

### 3.2 Performance on playback sounds in Iriomote

To evaluate the applicability of our model as an early warning system, we evaluated the performance of our model on the playback sounds in Iriomote Island, which is about 30 km from Ishigaki. Briefly, recordings from Ishigaki were replayed and re-recorded in Iriomote (more details in Kimura et al. (2025) [18]). Our model achieved high *F*_1_ scores for both species (0.76 for *P. leucomystax*, and 0.99 for *R. marina*) after adjusting the threshold for the Iriomote recordings. Without threshold adjustement, our detector exhibited an *F*_1_ score of 0.54 and 0.92 for *P. leucomystax* and *R. marina* respectively. The low *F*_1_ score for *P. leucomystax* at the original threshold (t=0.62) is attributed to low recall (0.38), as precision was sufficiently high (0.94) and comparable to the precision at the Ishigaki sites (0.93).

Subsequently, we investigated the performance of our detector in each individual recording session in Iriomote (Figure 3 and Table 4). Our lightweight model successfully detected nine out of eleven playback sessions for *P. leucomystax* (Table 4). Using the lower threshold optimized for Iriomote (t=0.42) resulted in a higher overall number of detections, but still missed the sessions on the 29th of July and the 28th of December. Detection of *R. marina* was successful for all playback sessions, while using the lower threshold (t=0.54) increased the total detections from 168 to 193 (out of 196).

**Fig. 3.**
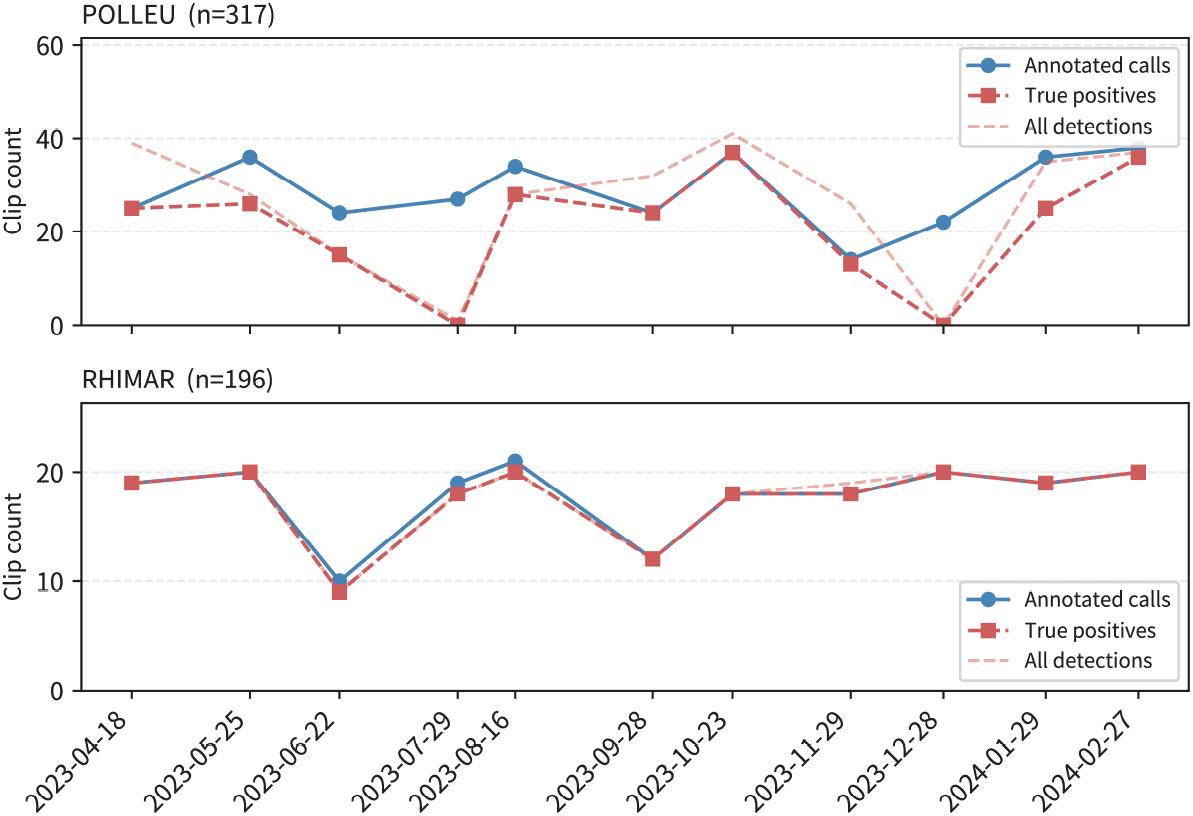
Performance of our lightweight model in playback recordings in Iriomote island, shown per individual recording session

### 3.3 Performance on an Australian cane-toad dataset

To further evaluate the performance of our lightweight model in a global context, we tested its performance on a cane toad dataset (*R. marina*) from Australia [23]. As the only common species between our 8-species classifier and the Australian cane toad dataset was *R. marina*, we grouped all other species into a single group (independently for each dataset) and focused on the performance of our detector solely on *R. marina*. Note, we conducted this test as a zero-shot evaluation, without any additional training on the Australian data, as our model was trained only with the data from Ishigaki island, Japan.

From a total of 22,006 positive samples in the cane toad dataset, 20,605 were successfully identified as *R. marina* at the Ishigaki-based threshold, achieving an *F*_1_ score of 0.89. The best *F*_1_ score, found by searching through a range of threshold values, was also 0.89, and achieved at a threshold of 0.77, with precision and recall showing only minor changes due to the different threshold (Table 5).

To confirm that our detector learned meaningful features specific to *R. marina* we conducted two ablation experiments. Firstly, we created a model of the same architecture as our lightweight detector but trained only the head classifier while keeping the weights of the convolutional features random. This variant shows how much the learned convolutional features contribute to performance. In other words, it shows how well a trained classifier head can perform on top of random, untrained features. Secondly, we created a model in which all weights, including both the convolutional features and the classification head, were set to random values, serving as a baseline that rules out artifacts of the architecture itself.

Figure 4 shows the confusion matrices of the three different models. Our fully trained model could successfully detect 20,605 positive samples. The model with the random convolutional features could only detect 6,867 positive samples, while the model with all random features could not detect any positive sample. These results confirm that our detector learned meaningful convolutional features during training that are highly specific to *R. marina*, and generalize successfully from the local Ishigaki recordings to the Australian dataset.

**Fig. 4.**
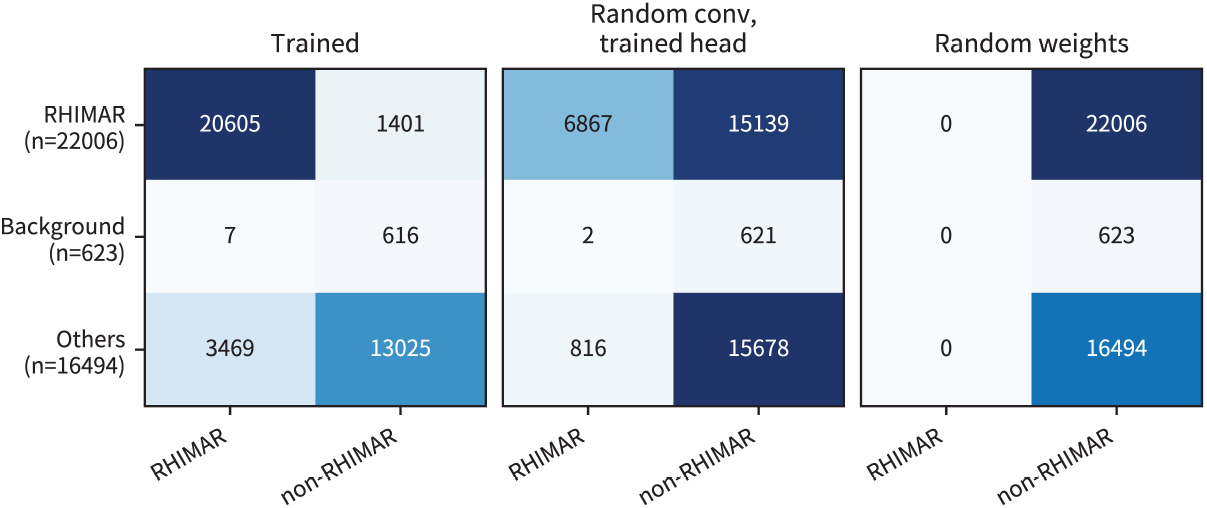
Performance of our lightweight model and two ablated variations in the cane toad dataset from Australia. Three confusion matrices are shown. **Trained:** Our lightweight detector, fully trained. **Random conv, trained head:** Our lightweight detector with random convolutional features and a trained head classifier. **Random weights**: Our lightweight detector with all weights (convolutional and head) set to random. The first two models use their own threshold values, computed independently from their respective validation datasets (Ishigaki data). The values shown in the confusion matrices represent the 0-shot performance of the trained and ablated models on the Australian dataset. The last model has no training and uses the same threshold as the first model (0.77)

### 3.4 Comparison to larger architectures

We compared our lightweight model against two larger architectures in three different variations, across all three evaluation datasets. We used EfficientNet-B0 trained from scratch, a half-width EfficientNet-B0 trained from scratch, and ResNet50 pre-trained on ImageNet. Table 6 shows the number of parameters and size of each model in its quantized state.

To isolate each architecture’s discriminative capacity from threshold calibration effects, we evaluated each model at its *F*_1_-optimal threshold per dataset. For the Ishigaki test set, thresholds were derived from the validation split. For the zero-shot evaluations on Iriomote and Australia, we report each model’s best achievable *F*_1_, removing threshold transfer as a confounding factor.

The proposed model achieved *F*_1_ scores that are competitive to the larger architectures, across all evaluation contexts (Table 7). On the Ishigaki test set, it outperformed all baselines for both invasive species (POLLEU *F*_1_ = 0.88, RHIMAR *F*_1_ = 0.87). On the zero-shot Iriomote evaluation, our proposed model achieved *F*_1_ = 0.76 for *P. leucomystax*. The best *F*_1_ score for *P. leucomystax* in Iriomote was achieved by ResNet50 pretrained on ImageNet (0.84). The other two variants performed similarly to our detector. For *R. marina* in Iriomote and Australia, our detector was the best performing model.

### 3.5 Inference time on low-power microcontrollers

Because our lightweight detector is intended for early-warning systems on remote islands, we used Edge Impulse to benchmark it across several common low-power MCU platforms, focusing on three metrics: inference time, RAM usage, and flash (ROM) usage. Edge Impulse reported that our model requires approximately 133 KB of flash (ROM) and 403 KB of RAM when deployed. The 133 KB flash footprint, which includes the 66.6 KB quantized model file plus the runtime code and buffers needed to execute it, fits comfortably on all platforms tested. RAM requirements, however, proved more restrictive as 403 KB exceeds the on-chip RAM of three of the devices evaluated: the Arduino Nano 33 BLE, the RP2040, and the nRF54L15. Although Edge Impulse returns inference-time estimates for these platforms, they cannot, in fact, run the model.

Among the platforms with sufficient RAM, inference time varied substantially (Table 8). The STM32N6, equipped with a dedicated neural-processing unit, completed inference in 4 ms. The Portenta H7 (84 ms) left ample margin for mel-spectrogram generation within the 3-second analysis window, as did the Sony Spresense (414 ms). These estimates suggest real-time operation is feasible on these platforms, though the actual on-device latency depends strongly on implementation factors such as memory placement and compiler optimization, and must be confirmed by direct measurement (future work).

## 4 Discussion

In this paper, we presented a lightweight deep-learning model for the early detection of two invasive frog species, *R. marina* and *P. leucomystax*. Our model was trained with audio samples recorded in Ishigaki island, Japan [18]. In addition to the held-out test dataset from Ishigaki, we evaluated our detector using data from two additional locations: playback sounds from Iriomote Island, Japan (30 km from Ishigaki), and an Australian cane toad dataset. Overall, our results indicate that lightweight detectors can be successfully utilized for the detection of *R. marina* and *P. leucomystax*, with performance comparable to that of larger architectures that require more power-hungry infrastructure.

On the held-out Ishigaki test set, the detector performed strongly for both invasive species, achieving an *F*_1_ score of 0.88 for *P. leucomystax* and 0.87 for *R. marina*, with high precision for both species (0.93 and 0.88 respectively) (Table 2). For the Iriomote and Australia datasets we report two threshold values: the original threshold that maximized the score on Ishigaki, and the threshold that maximized it at the respective location. At Iriomote, we found that *R. marina* could be reliably detected even with the original Ishigaki-based threshold (*F*_1_ = 0.92), while *P. leucomystax* required a lower, locally optimized threshold to achieve an *F*_1_ score of 0.76. In the Australian cane toad dataset, our detector achieved high performance even with the Ishigaki-based threshold (*F*_1_ = 0.89).

The sensitivity of deep-learning detectors to the choice of threshold is well documented in the bioacoustics literature, with optimal values varying both among species [34, 35] and among recording sites due to differences in background noise and recording conditions [36]. As shown by our analysis of the Iriomote playback recordings (Table 3), our lightweight detector is not immune to this problem, as *P. leucomystax* suffered from a low recall rate at the Ishigaki-based threshold (while precision was kept high). However, the fact that simply reducing the threshold for Iriomote brings performance close to that of the Ishigaki test set (*F*_1_ = 0.76 in Iriomote, *F*_1_ = 0.88 in Ishigaki) suggests that our model can still discriminate *P. leucomystax*, but it does so with a lower confidence.

**Table 3.** Cross-site evaluation on the Iriomote playback recordings, for POLLEU (*P. leucomystax* ) and RHIMAR (*R. marina*). TP/FP/FN = counts of true positives, false positives, and false negatives over 3-second windows.

| Species | Strategy | TP | FP | FN | F1 | Precision | Recall | Threshold |
| --- | --- | --- | --- | --- | --- | --- | --- | --- |
| POLLEU | Ishigaki threshold | 119 | 8 | 198 | 0.54 | 0.94 | 0.38 | 0.62 |
| POLLEU | Iriomote-optimal | 229 | 53 | 88 | 0.76 | 0.81 | 0.72 | 0.42 |
| RHIMAR | Ishigaki threshold | 168 | 0 | 28 | 0.92 | 1.00 | 0.86 | 0.79 |
| RHIMAR | Iriomote-optimal | 193 | 1 | 3 | 0.99 | 0.99 | 0.98 | 0.54 |

**Table 4.** Detection of invasive species on Iriomote playback recordings by session date. Pos. = total annotated positive windows per night. For each species, results are shown at the original Ishigaki-optimized threshold and at the *F*_1_-optimal threshold on the Iriomote set itself. Shaded cells indicate zero detections (at either threshold).

| Date | *P. leucomystax* |  |  | *R. marina* |  |  |
| --- | --- | --- | --- | --- | --- | --- |
| | Pos. | $t=0.62$ | $t=0.42$ | Pos. | $t=0.79$ | $t=0.54$ |
| 2023-04-18 | 25 | 19 | 25 | 19 | 18 | 19 |
| 2023-05-25 | 36 | 2 | 26 | 20 | 18 | 20 |
| 2023-06-22 | 24 | 5 | 15 | 10 | 7 | 9 |
| 2023-07-29 | 27 | 0 | 0 | 19 | 15 | 18 |
| 2023-08-16 | 34 | 3 | 28 | 21 | 15 | 20 |
| 2023-09-28 | 24 | 23 | 24 | 12 | 12 | 12 |
| 2023-10-23 | 37 | 25 | 37 | 18 | 18 | 18 |
| 2023-11-29 | 14 | 7 | 13 | 18 | 17 | 18 |
| 2023-12-28 | 22 | 0 | 0 | 20 | 13 | 20 |
| 2024-01-29 | 36 | 11 | 25 | 19 | 19 | 19 |
| 2024-02-27 | 38 | 24 | 36 | 20 | 16 | 20 |
| **Total** | **317** | **119** | **229** | **196** | **168** | **193** |

Still, despite the increase in the *F*_1_ score after threshold reduction, our detector completely missed two of the eleven sessions in the Iriomote playback recordings (29th July and 28th December). It is worth noting that, in the original study of Kimura et al. [18], BirdNET also exhibited a lower number of detections on 29th July and 28th December, highlighting the general problem of cross-site generalization even for large-scale foundational models.

The findings from our threshold analysis translate into concrete deployment guidance for biodiversity monitoring. When a system with a deep-learning detector becomes operational at a new site, it is recommended to conduct initial trials with playback recordings to allow for calibration of the threshold value. In the absence of such calibration data, the threshold can be lowered as a precaution, though this would increase the false-positive rate and, in turn, the manual validation effort required. Finally, if confidence scores remain low even after threshold adjustment, retraining with local or augmented data may be needed to reach acceptable performance. However, not all species may exhibit reduced performance at new sites, as our detector could reliably detect *R. marina* in Iriomote, and even in the dataset from Australia (Table 3 and Table 5).

**Table 5.**
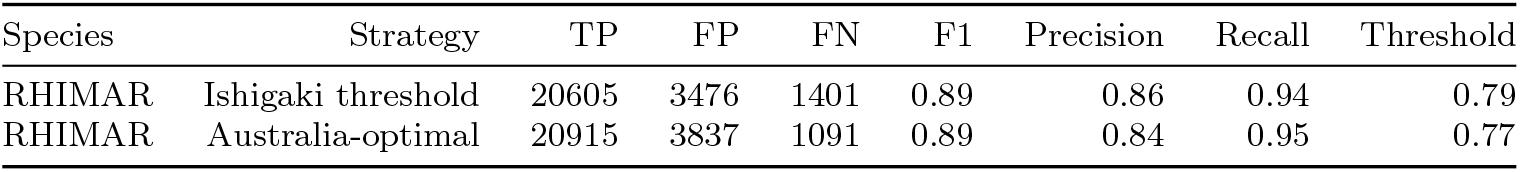
Evaluation of our lightweight detector model on the Australian Cane Toad dataset (RHIMAR = *R. marina*). TP/FP/FN = counts of true positives, false positives, and false negatives over 3-second windows.

**Table 6.**
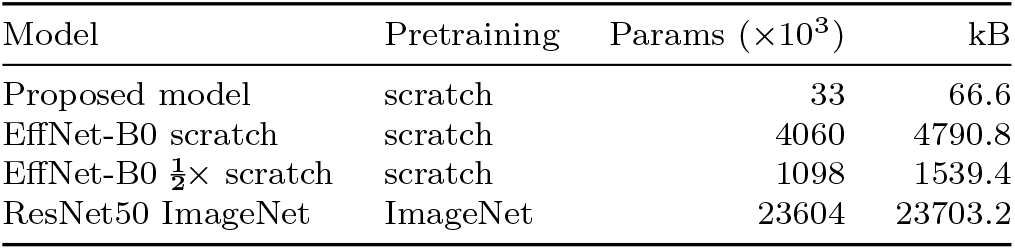
Model size and parameter counts for the proposed lightweight detector compared against the fine-tuning baselines (all metrics computed from the INT8 model). The Pretraining column indicates whether the backbone was initialized from ImageNet weights or trained from scratch.

We also compared our lightweight detector against three larger baseline architectures (Table 7). Across all different evaluation contexts (Ishigaki, Iriomote, Australia), our detector was among the highest-performing models despite its much smaller size. It should be noted, however, that because each model was trained and evaluated only once, we do not claim that our detector is the best-performing model with absolute certainty. Rather, our results evidence that our lightweight detector can achieve detection performance comparable to that of architectures with orders of magnitude more parameters.

**Table 7.** F1 score for the proposed model against three standard baselines across all evaluation contexts (all metrics computed from the saved INT8 model). Species: POLLEU = *P. leucomystax*, RHIMAR = *R. marina*. Sites: ISH = Ishigaki test set, IRI = Iriomote playback, AUS = Australian Cane Toad dataset. Cells from the Ishigaki test set use each model’s validation-derived F1-optimal threshold. The two additional site columns (IRI, AUS) use each model’s F1-optimal threshold on the respective evaluation set itself.

| Model | POLLEU (ISH) | RHIMAR (ISH) | POLLEU (IRI) | RHIMAR (IRI) | RHIMAR (AUS) |
| --- | --- | --- | --- | --- | --- |
| Proposed model | 0.88 | 0.87 | 0.76 | 0.99 | 0.89 |
| EffNet-B0 scratch | 0.86 | 0.67 | 0.77 | 0.85 | 0.86 |
| EffNet-B0 $\frac{1}{2}\times$ scratch | 0.74 | 0.82 | 0.76 | 0.91 | 0.87 |
| ResNet50 ImageNet | 0.84 | 0.63 | 0.84 | 0.96 | 0.78 |

Our lightweight detector is expected to run on several low-cost MCUs, with inference time ranging from 4 ms on the STM32N6 to 414 ms on the Sony Spresense (Table 8). The feasibility of running our detector on a diverse range of low-power MCUs offers a concrete advantage for invasive species monitoring. Instead of relying on SD cards that must be manually retrieved, an MCU-based system can perform real-time detection on the device itself, enabling rapid response by on-the-ground eradication teams at the earliest stages of an invasion.

**Table 8.** Edge Impulse–estimated inference time and on-device feasibility for the profiled platforms (ordered as in Table 1). Feasibility is set by on-chip RAM: the model’s 403 KB activation arena exceeds the RAM of the three most constrained boards, which therefore cannot run the model despite Edge Impulse reporting a latency for them (its estimate is independent of the memory-fit check). Inference time excludes mel-spectrogram generation; the analysis window is 3 s. The STM32N6’s 4 ms reflects its integrated NPU.

| Device | Estimated Inference (ms) | Runs on-device |
| --- | --- | --- |
| Arduino Nano 33 BLE | 896 | No (insufficient RAM) |
| Nordic nRF54L15 | 383 | No (insufficient RAM) |
| Raspberry Pi RP2040 | 2,030 | No (insufficient RAM) |
| Portenta H7 | 84 | Yes |
| Sony Spresense | 414 | Yes |
| STM32N6570-DK | 4 | Yes |

Profiling with Edge Impulse was critical for determining which devices can actually run our lightweight model. While the flash (ROM) required to store the model fits within the available memory of all the low-power devices considered, the RAM required at inference time, dominated by the intermediate activations, exceeded the on-chip RAM of three MCUs, precluding their use with our model.

An MCU-based detector is best understood as a triage system, rather than a replacement for expert verification: it can continuously screen the soundscape and flag candidate detections, which then trigger targeted human follow-up. This could be a field visit, or, if possible, online retrieval of the flagged audio segment. In this role, a modest false-positive rate is acceptable, since each flag prompts a single verification, whereas a missed invasion poses a greater ecological risk. The principal practical advantage of running inference on an MCU is power: drawing substantially lower power than a single-board computer, an MCU can achieve comparable monitoring autonomy from a much smaller battery and solar panel, easing deployment and servicing at remote sites. The final power consumption will be determined by the MCU and any additional peripherals, such as storage or wireless transmission.

Through on-device classification, only sporadic detection events need to be transmitted, rather than continuous audio. As a result, invasive species monitoring with MCUs can use low-bandwidth, long-range protocols such as LoRa or satellite IoT that remain available even where WiFi or cellular coverage does not exist. This reduced dependence on high-bandwidth connectivity broadens the locations where autonomous acoustic monitoring can be deployed, such as deep within forests or in the remote interiors of Natural World Heritage sites. Moreover, even where conventional coverage exists, transmitting only detection events reduces power consumption, bandwidth usage, and data transmission costs.

A further advantage of on-device inference concerns data privacy. Traditional acoustic monitoring that saves raw audio to SD cards will inevitably capture any nearby human conversations, raising privacy concerns about the handling and storage of the raw audio data. Utilizing lightweight detectors directly on the device negates the need for long-term storage of raw audio and enables the transmission/storage of detection events only. It thus avoids storing potentially sensitive audio, an increasingly relevant consideration for acoustic monitoring near populated areas.

Our current analysis focused on the performance of the lightweight detector to demonstrate its feasibility and the limitations of its detection capacity. Our future work will include developing a prototype to verify operation with low-power MCUs, troubleshoot the long-term, real-world operation, and power measurements for the entire system.

## 5 Data availability

The trained detector and code to reproduce the results of this paper can be found in https://github.com/deligkarisk/lightweight detector anurans release

## 6 Acknowledgments

We are grateful to Kimura Kaede for helpful comments on an earlier version of the manuscript. We are also grateful for the help and support provided by the Scientific Computing and Data Analysis Section and the Marine and Terrestrial Sciences Section of the Research Support Division at OIST.

## 8 Author information

Conceptualization: KD, GS; Methodology and software: KD; Formal analysis: KD, MST; Writing, original draft: KD; Writing, review and editing: KD, MST, GS; All authors read and approved the final version of the manuscript.

